# Restoring statistical validity in group analyses of motion-corrupted MRI data

**DOI:** 10.1101/2021.06.15.448467

**Authors:** Antoine Lutti, Nadège Corbin, John Ashburner, Gabriel Ziegler, Bogdan Draganski, Christophe Phillips, Ferath Kherif, Martina F. Callaghan, Giulia Di Domenicantonio

## Abstract

Motion during the acquisition of magnetic resonance imaging (MRI) data degrades image quality, hindering our capacity to characterize disease in patient populations. Quality control procedures allow the exclusion of the most affected images from analysis. However, the criterion for exclusion is difficult to determine objectively and exclusion can lead to a suboptimal compromise between image quality and sample size. We provide an alternative, data-driven solution that assigns weights to each image, computed from an index of image quality using restricted maximum likelihood. We illustrate this method through the analysis of brain MRI data. The proposed method restores the validity of statistical tests, and performs near optimally in all brain regions, despite local effects of head motion. This method is amenable to the analysis of a broad type of MRI data and can accommodate any measure of image quality.

## Main

Movement notoriously degrades magnetic resonance imaging (MRI) data, leading to prolonged examinations and increased costs in clinical applications ^1,2^. Head movement also impacts the estimates of brain features extracted from MRI data ^3–7^, and can lead to spurious detection or suppression of brain change in neuroscience studies ^2^. This issue is particularly acute for the study of non-compliant patient populations, where the effects of brain disease and head motion cannot be separated ^8,9^. Quality control procedures exist that help mitigate the effect of image degradation on analysis results. These procedures require an assessment of data quality, provided by a motion degradation index (MDI) ^10–16^. MDIs computed using dedicated image analysis routines require little labour investment and have become instrumental in the oversight of the large data cohorts that have emerged in recent years ^17–21^. Recent findings suggest that they may provide higher sensitivity for detecting motion artefacts than visual assessment ^13,22^ and, combined with supervised learning methods ^14,23,24^, might allow the automated identification of images usable in subsequent analyses.

Quality control is typically followed by dichotomising the data into images that are either suitable (“accept”) or unsuitable (“exclude”) for analysis. Here, the threshold value of the MDI between the “exclude” and “accept” categories can be difficult to determine. Also, the effects of motion on MRI data are continuous ^13,22^ and it is likely that no hard categorization of the data might achieve optimal compromise between image quality and sample size. We propose an alternative method that assigns a weight to each image within a cohort, computed from its MDI value using the restricted maximum likelihood (REML) algorithm ^25^. The weights are specific to each image and down-weight low quality images in subsequent analyses. We illustrate this method through the analysis of a large cohort (1,432 participants) of quantitative MRI (qMRI) data. qMRI data are *in vivo* biomarkers of brain microstructure^26–28^ that have great potential for clinical neuroscience ^29–31^. We use the MDI introduced in Castella *et al.*^16^, a validated marker of motion degradation.

We show that in conventional analyses, degradation of image quality due to motion invalidates any assumption of identical variance for all samples (‘homoscedasticity’). While heteroscedasticity has little impact on the model coefficient estimates in a general linear model, the standard error of the coefficients can be poorly estimated, leading to invalid statistical inference^32^. The proposed method, called QUIQI for ‘analysis of QUantitative Imaging data using a Quality Index’, restores homoscedasticity, ensuring the validity of statistical tests. This global approach provides near optimal results in whole-brain analysis of neuroimaging data, despite local effects of motion. The framework has been implemented in the popular, open-source neuroimaging analysis software Statistical Parametric Mapping (www.fil.ion.ucl.ac.uk/spm, Wellcome Centre for Human Neuroimaging). The framework is flexible and amenable to other MDIs and to the analysis of other types of MRI data.

## Results

### QUIQI is integrated in the analysis of the MRI data

We illustrate this method on the group-level analysis of quantitative maps of the MRI parameter R2*. This parameter is sensitive to iron and myelin density and is computed from acquired MRI data by fitting a mono-exponential decay model (Fig. 1a). A previously validated MDI is computed from these maps, as described in Castella et al. ^16^ (Fig. 1b). With QUIQI, the values of the MDI – specific for each image sample – are inserted into the REML algorithm in the form of basis functions that capture the relationship between image noise and the MDI. From the empirical demonstration of this relationship in Fig. 2, the basis functions are comprised of multiple powers of the MDI. REML computes a noise covariance matrix *V* that captures the noise level in each image. The matrix *V* is diagonal because noise is uncorrelated between images, i.e., participants (Fig. 1c). In a final step, image-specific weights are computed from the noise covariance matrix and the parameters of the general linear model are estimated voxel-wise, resulting in a weighted least-square analysis (WLS, Fig. 1d).

**Fig. 1.**
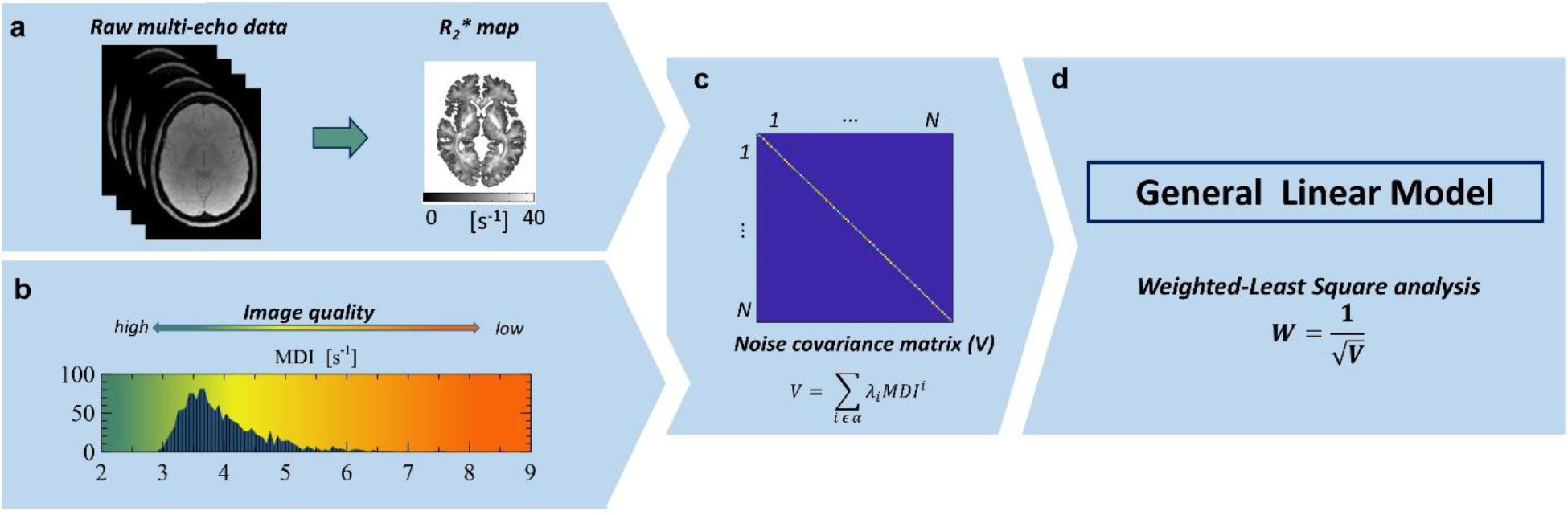
QUIQI integrates correction of motion degradation into the analysis of MRI data. For the current application, analysis data are quantitative maps of the MRI parameter R2* (**a**). QUIQI requires a value of the MDI for each data set of the analysis. Here, we show the distribution of the MDI values across the images used for analysis (N=1,432) (**b**). With QUIQI, basis functions are computed from powers of the MDI and inserted into REML ^25^ for the computation of the noise covariance matrix *V* (**c**). The set of powers of the MDI, α, is pre-defined by the user. From *V*, weights are computed that are used in the general linear model for data analysis (**d**).

**Fig. 2.**
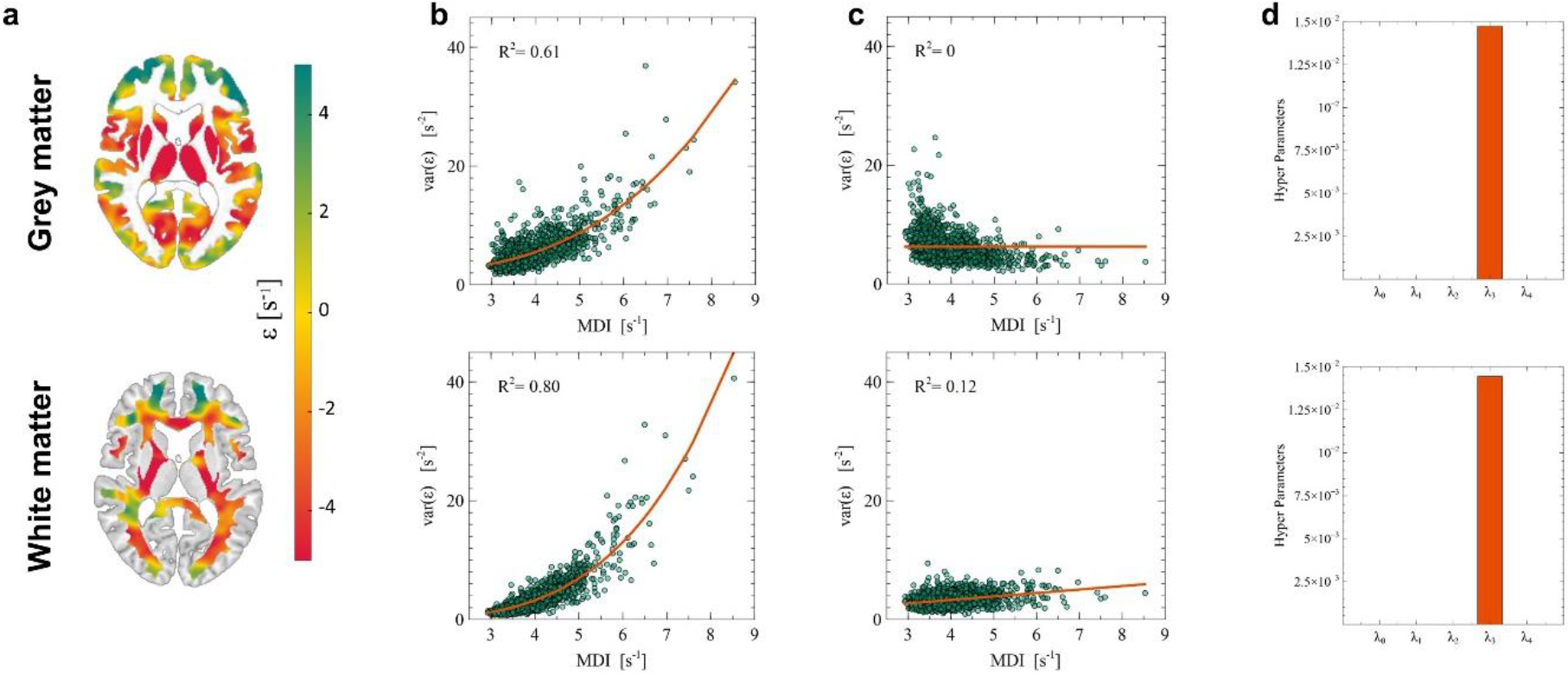
QUIQI restores homoscedasticity of the residual noise distribution in an analysis. Following model fitting, maps of the residuals are computed for each individual image (example shown in **a**). An estimate of image noise is computed from the variance across these residual maps and plotted against the MDI for OLS (**b**) and WLS analyses (**c**). Enforcing positivity for the hyper-parameters (λ_i_>0) leads to a dominant cubic power in the modelling of image noise by REML (**d**).

### The motion degradation index is a predictor of residual noise

The use of REML relies on the empirical observation that in a conventional analysis (ordinary least squares, OLS), noise in MRI data can be accurately modelled from the MDI. We computed an estimate of image noise as the spatial variance of the residual maps in each sample image (*var(ε)*). Fig. 2a shows an example of such residual maps for grey and white matter. In an OLS analysis, the polynomial dependence of residual noise on the MDI highlights heteroscedasticity in the data, which leads to misestimation of the precision of parameter estimates and undermines the validity of statistical tests (Fig. 2b). From this dependence, QUIQI uses powers of the MDI as basis functions for estimating the noise covariance matrix. The resulting residual noise is independent of the MDI, restoring the homoscedasticity of the data (Fig. 2c). The REML hyper-parameter estimates (λ_i_), obtained with a positivity constraint, illustrate the dominant contribution of the cubic power of the MDI (MDI^3^) to the noise covariance matrix, consistently for grey and white matter (Fig. 2d).

We conducted Engle’s ARCH tests of residual heteroscedasticity in each voxel of the MRI data. In grey matter, the null hypothesis of no ARCH effects could be rejected in 82% and 12% of voxels for OLS and WLS respectively (p<0.05, see Extended Data Fig. 1). For the WLS analysis, these voxels were mainly located in sub-cortical regions, regions affected by magnetic field inhomogeneities, or at the interface of brain tissue with its surroundings. In white matter, the null hypothesis could be rejected in 92% and 9% of voxels for OLS and WLS respectively (p<0.05). Incidentally, WLS also reduced the number of voxels where the hypothesis of standard normally distributed noise could be rejected (Kolmogorov-Smirnov tests).

### Optimal modelling of noise from the MDI

The evidence lower bound (ELBO), or negative variational free energy, is a model selection criterion that favours reducing residual errors while also penalizing model complexity. The optimal noise model computed from the MDI maximised the ELBO provided by REML. At the global level of a given tissue type, i.e., grey or white matter, a gain in ELBO of up to two orders of magnitude was obtained with WLS compared to OLS analyses (Fig. 3a). Consistent with Fig. 2d, the maximum gain was obtained using the MDI cubed (i.e., MDI^3^) as a basis function in the REML estimation (*global optimal model*). Including additional powers of the MDI did not increase the ELBO.

**Fig. 3.**
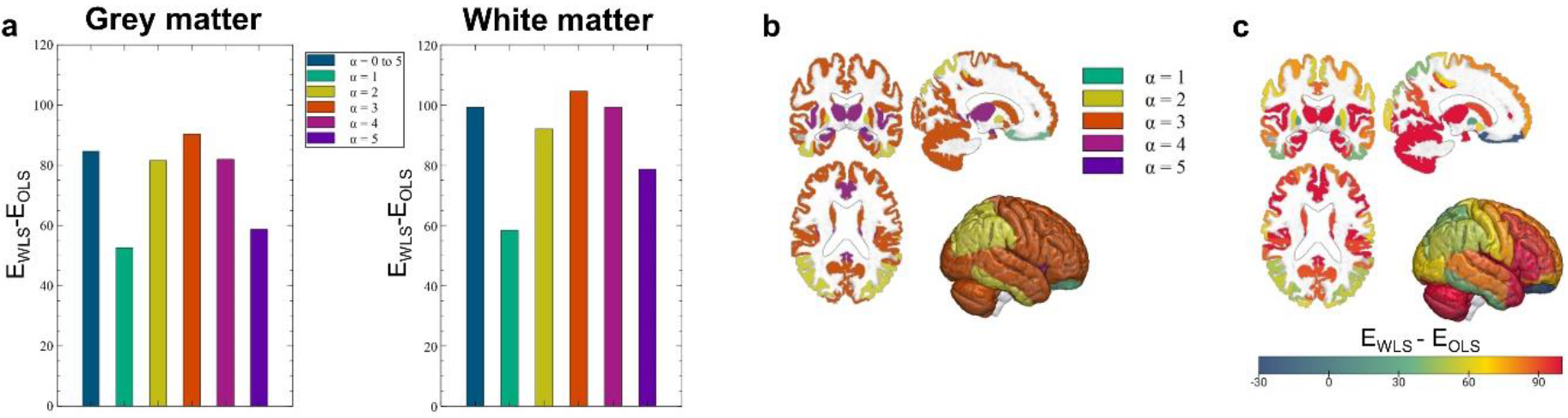
Global and local analysis of the REML ELBO. At the global level of a whole tissue type (grey or white matter), the gain in ELBO compared to OLS analyses is maximal with a single basis function in the REML estimation that contains the cubic power of the MDI values (α=3, **a**). In grey matter regional analyses, the global optimal model (MDI^3^) was also the local optimal within most regions, primarily located in frontal areas (**b**). In posterior regions, the local optimal model involved the square power of the MDI. With the global optimal model, the local gain in ELBO showed a gradient in the anteroposterior direction, with the highest gain in frontal areas (**c**). This is consistent with typical motion of study participants during MRI examinations.

QUIQI is primarily intended for the analysis of entire images. However, for the purpose of assessing QUIQI’s ability to correct for local degradation of image quality due to head motion, we repeated the analysis separately for each region of a grey matter atlas. The global optimal model (MDI^3^) led to the highest gain in ELBO in 68% of regions. In the remaining regions, the ELBO from the global optimal model was smaller than its local counterpart by an average of 5.2. Although substantial in terms of model evidence, these differences are small compared to the gain over OLS analyses. Regions where the global and local optimal models were identical were located primarily in frontal areas, while posterior areas tended to exhibit locally optimal models with a lower power of the MDI (Fig. 3b). This anterior-posterior gradient is also apparent in the increase in ELBO compared to OLS analyses: the highest gains are observed in frontal regions (Fig. 3c).

### QUIQI increases analysis sensitivity to brain differences

To illustrate the effect of QUIQI on the detection of brain-related differences in neuroscience, we conducted statistical F-tests of the dependence of the MRI data on age. As previously reported ^33,34^, the most prominent age-related differences in R2* were located in sub-cortical grey matter due to a local increase in iron concentration with age and in frontal white matter due to a peak in axonal myelination around midlife ^35^ (Fig. 4a).

**Fig. 4.**
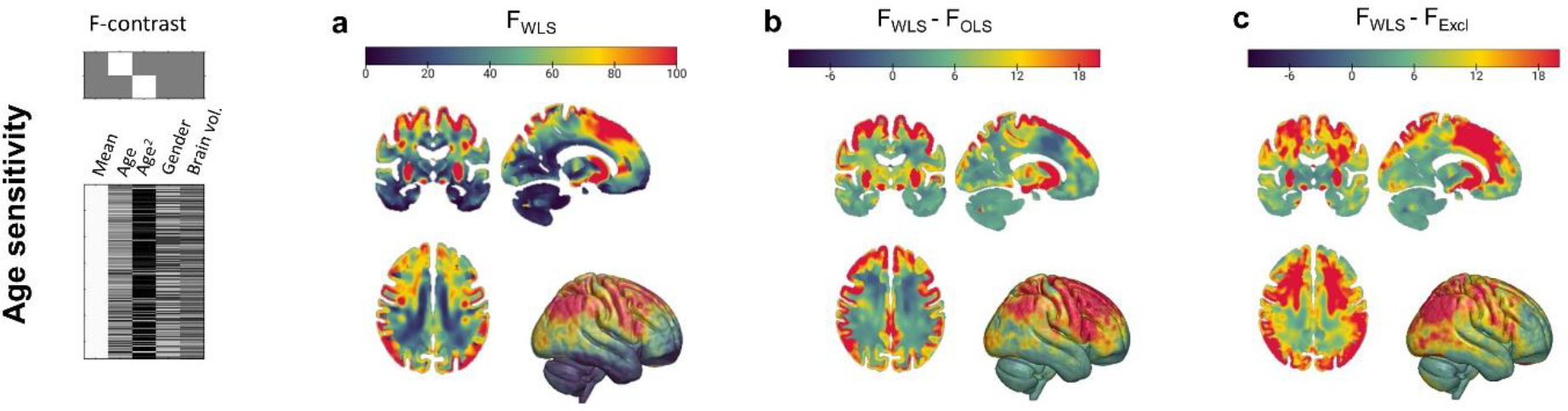
QUIQI increases the sensitivity of MRI data analysis. The effect of QUIQI on the analysis of brain-related differences was assessed using statistical F-tests of the dependence of the MRI data with age (**a**). Compared with OLS analyses, QUIQI leads to region-specific decreases or increases in F-scores due to the restored noise homoscedasticity (**b**). Exclusion of the 30% most degraded images was required to restore noise homoscedasticity in OLS analyses. WLS analyses yield higher age-sensitivity over the whole brain (**c**).

Compared with OLS analyses, QUIQI leads to region-specific decreases or increases in F-scores (Fig. 4b) but because QUIQI restores noise homoscedasticity, WLS analyses are more sensitive to true age effects. We compared the age sensitivity of WLS analyses with that of OLS analyses after exclusion of the fraction of the images with the highest MDI values (i.e., the most degraded images, see Extended Data Fig. 2a). From the multiple fractions considered (3, 7, 13, 20 and 30% ^12,14,15,23^), we selected the one that led to similar noise homoscedasticity to WLS analyses. In grey matter, this was achieved after removing 30% of the images (R^2^=0.11, see Extended Data Fig. 2b). With this fraction of excluded data, heteroscedasticity was still present in white matter (R^2^=0.43) but higher values were deemed too prohibitive to be considered. WLS analyses led to higher age-sensitivity than exclusion OLS analyses in both tissue types (Fig. 4c).

### QUIQI preserves the specificity of statistical analyses

We verified the specificity of the proposed technique by monitoring the rate of false positives in statistical tests. In the first test, the age regressor was scrambled using random permutations, so that any test indicating age-related difference would be considered a false positive. Results (Table 1, Line 1) showed that the false positive rate was within the expected range at the voxel and cluster levels, for both OLS and WLS analyses (p<0.05, FWE-corrected; Table 1). In the second test, a sub-cohort was selected within a narrow age range and group comparisons were conducted within this sub-cohort, so that group differences would be considered as false positives. Results (Table 1, Line 2) showed that the false positive rate increased similarly for OLS and WLS analyses in unbalanced group comparisons ^36^. For white matter, a false positive rate within 0.05 was found for cluster-level inference with N_1_ ≈ 5 or more images in the first group. More false positives were observed for grey matter. The false positives were primarily located in cortical regions affected by magnetic field inhomogeneities (e.g. orbitofrontal cortex and temporal lobes, see Extended Data Fig. 3).

**Table 1.** Specificity of MRI analyses: false positive rates in one-sample and two-sample analyses. The false positive rates are reported at the voxel/cluster-levels. For one-sample t-tests, the false positive rate was below 0.05, the p-value used for statistical significance, for both OLS and WLS analyses. For two-sample t-tests the false positive rate, reported for different numbers of subjects N_1_ in the first group, increased similarly for OLS and WLS analyses in unbalanced group comparisons ^36^. The higher number of false positives in grey matter arises from cortical regions affected by magnetic field inhomogeneities leading to bias on the R2* estimates (see Extended Data Fig. 3)

|  |  | GM |  |  |  |  | WM |  |  |  |  |
| --- | --- | --- | --- | --- | --- | --- | --- | --- | --- | --- | --- |
| One sample t-tests | OLS | 0.02/0.022 |  |  |  |  | 0.029/0.023 |  |  |  |  |
|  | WLS | 0.027/0.022 |  |  |  |  | 0.026/0.013 |  |  |  |  |
| Two sample t-tests |  | N<sub>1</sub>=2 | N<sub>1</sub>=5 | N<sub>1</sub>=10 | N<sub>1</sub>=30 | N<sub>1</sub>=60 | N<sub>1</sub>=2 | N<sub>1</sub>=5 | N<sub>1</sub>=10 | N<sub>1</sub>=30 | N<sub>1</sub>=60 |
|  | OLS | 0.860<br>/0.245 | 0.509<br>/0.104 | 0.203<br>/0.055 | 0.074<br>/0.049 | 0.051<br>/0.042 | 0.618<br>/0.182 | 0.254<br>/0.066 | 0.132<br>/0.043 | 0.076<br>/0.021 | 0.072<br>/0.032 |
|  | WLS | 0.895<br>/0.268 | 0.601<br>/0.112 | 0.332<br>/0.061 | 0.102<br>/0.047 | 0.075<br>/0.046 | 0.592<br>/0.167 | 0.256<br>/0.05 | 0.117<br>/0.03 | 0.068<br>/0.024 | 0.071<br>/0.033 |

### Modelling motion-related variance from covariates in the analysis design does not restore homoscedasticity

As a potential alternative to QUIQI, we modelled motion-related variance in the data by inserting dedicated regressors in the design matrix ^37^. On the model of the REML basis functions, these regressors contained the 1st to 4th powers of the MDI values. For a subset of the data within a narrow age range, we conducted statistical F-tests of the variance of the R2* maps associated with these regressors. For OLS analyses, statistically significant results were found in 3.2% and 5.5% of voxels in grey and white matter respectively (p<0.05, FWE-corrected). For grey matter, these voxels were primarily located in frontal regions (see Fig. 5a). For WLS analyses, statistically significant results were only found in 0.1% of voxels in grey and white matter (p<0.05, FWE-corrected). Residual analysis showed that noise heteroscedasticity remains in the data for both tissue types with OLS analyses, despite the motion regressors. Noise heteroscedasticity is not present in WLS analyses (Fig. 5b).

**Fig. 5.**
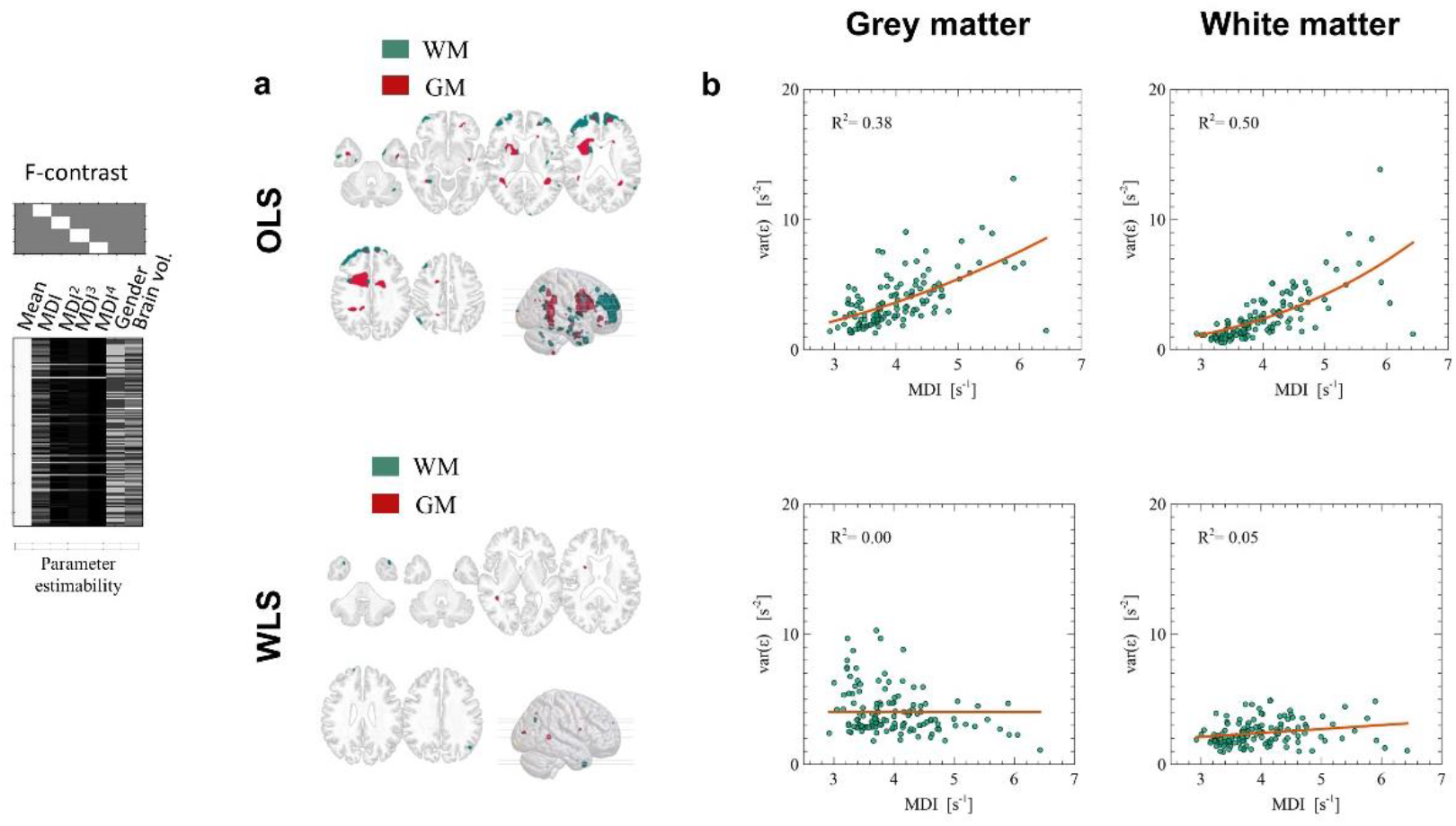
Using motion regressors as design covariates does not restore heteroscedasticity. In OLS analyses, statistical F-tests of the variance of the R2* maps associated with powers of the MDI show statistically significant results in 3.2% and 5.5% of voxels in grey and white matter respectively (p<0.05, FWE-corrected) (**a**). In WLS analyses, significant results are found in 0.1% of voxels only (p<0.05, FWE-corrected). Noise heteroscedasticity remains present for both tissue types in OLS analyses, despite the motion regressors, but is not present in WLS analyses (**b**).

## Discussion

In this study, we introduce a method that accounts for the degradation of image quality due to motion in the analysis of MRI data. This method addresses an important limitation of existing approaches for quality control, which only enable the removal of the most degraded images from analysis. From an index of image quality, the proposed method computes weights that capture the noise level in each image, leading to increased sensitivity to brain change in an analysis. This method is based on restricted maximum likelihood, available in most image analysis software suites. The presented implementation within the SPM software ^38^ is commonly used to account for differential noise levels in e.g. group-comparison studies (‘*non-sphericity’*). Here, we extend this methodology to the statistical analysis of structural MRI data, using an index of image quality to estimate the noise level in each individual image. We validated this method using a large cohort of 1,432 subjects, which allowed the design of multiple analyses to test different aspects of the method.

In conventional analysis methods, the increase of the noise level due to motion leads to a violation of the homoscedasticity assumption of statistical tests. By estimating the noise level in each image from its MDI value, QUIQI restores the validity of this assumption, both at the global level of a tissue type (i.e. grey or white matter) and in individual voxels of the MRI data. Voxels where significant heteroscedasticity remain with QUIQI are primarily located in regions where other non-motion mechanisms may be at play: sub-cortical areas (delineation errors), interface of brain tissue (partial volume effects) and regions of inhomogeneous magnetic field (bias of the R2* estimates). We tested an alternative method to QUIQI, based on the insertion of the MDI as a covariate in the design matrix of the analyses. However, we show that this alternative method does not successfully correct for noise heteroscedasticity.

To illustrate the effect of QUIQI on the analysis of MRI data, we conducted analyses of age-associated brain differences. OLS analyses led to spurious statistical results due to noise heteroscedasticity in the data. Restoring noise homoscedasticity in OLS analyses required the removal of the 30% of the images most affected by motion in grey matter (‘exclusion analyses’). This ratio was even higher for white matter. The sensitivity of WLS analyses to age-related differences was superior to that of exclusion analyses over the whole brain. The higher sensitivity of WLS analyses did not inflate the rate of false positives (Table 1).

QUIQI corrects local effects of head motion optimally – or near optimally – in whole-brain analyses, the archetypal use of MR images for neuroscience. The increase in ELBO with QUIQI was largest in frontal brain regions and the local optimal noise model involved a higher power of the MDI there. This is consistent with the supine position of the study participants during MRI examination, with the back of the head resting on the scanner table, and provides further evidence of the ability of QUIQI to correct the effects of motion on MRI data quality.

We implemented QUIQI by enforcing positivity of the hyper-parameter values estimated by REML (spm_reml_sc). This is by no means a requirement because allowing negative hyper-parameter values works equally well (Extended Data Fig. 4). Our choice was primarily guided by the fact that enforcing positivity allowed us to identify a single basis function (MDI^3^) as sufficient to effectively model noise in the data. With a single basis function, local analysis results reflect the ability of the method to correct for local effects of motion in a global analysis of a whole tissue type (e.g. grey or white matter). This would not be the case with several basis functions because the hyper-parameter values, which combine the basis functions in the estimation of the noise covariance matrix, would differ in a local and global analysis. Here, we provide a version of spm_est_non_sphericity that calls spm_reml_sc to enforce positive hyperparameter estimates ^39^. We also provide a customized version of the hMRI toolbox^40^ with a dedicated QUIQI module available from the GUI ^41^. This module allows the plotting of image noise versus the MDI (‘‘QUIQI check’’) to help users identify the optimal set of basis functions for the REML estimation.

The proposed method is amenable to all types of MRI data and motion degradation indices. Here, we emphasised the principles of the method and provided a detailed assessment of its performance. For illustration, we therefore chose the analysis of brain phenotype data that can be computed from one raw image type, i.e., quantitative maps of the MRI parameter R2*. The extension of this method to the analysis of quantitative maps computed across multiple types of raw images, each with their own degree of motion degradation, is currently ongoing. Another important field of application is the analysis of differences in brain morphology (e.g. grey matter volume or cortical thickness), the most widespread phenotypical measures extracted from MR images. Such applications will highlight the effect of image processing (segmentation) on the sensitivity of analysis to motion. While QUIQI can be readily used with different MDIs, we highlight the importance of the specificity and sensitivity of the index to motion degradation^16^, which drive the efficacy of the method. Potential sources of confound on image-based MDIs (e.g. brain disease) should be closely investigated (Extended Data Note 1).

## Methods

### Participant cohort

MRI data was acquired on 1,432 healthy research participants (743 females), as part of ‘BrainLaus’ (https://www.colaus-psycolaus.ch/professionals/brainlaus/) ^42–44^, a nested project of the PsyCoLaus/CoLaus study ^45,46^. The distribution of the motion degradation index values and participants’ ages are shown in Extended Data Fig. 5. The dependence of the index on the participants’ age is also shown there.

### MRI acquisition, pre-processing and analysis

#### MRI acquisition

The MRI protocol consisted of three multi‐echo 3D fast low angle shot (FLASH) acquisitions with magnetization transfer (MTw), proton density (PDw) and T1 (T1w)‐ weighted contrasts respectively. The repetition time and nominal flip angle were 24.5ms/6°, 24.5ms/6° and 24.5ms/21° respectively. The MTw contrast was achieved with a Gaussian‐shaped RF pulse prior to the excitation (4 ms duration, 220° nominal flip angle, 2 kHz frequency offset from water resonance). 6/8/8 echo images were acquired for the MTw/PDw/T1w contrasts with a minimal echo time of 2.34 ms and an inter-echo spacing of 2.34 ms. The image resolution was 1 mm^3^ isotropic, the field of view was 256 × 240 × 176. Parallel imaging was used along the phase‐encoding direction (acceleration factor 2; GRAPPA reconstruction ^47^). Partial Fourier was used in the partition direction with acceleration factor 6/8. The acquisition time was 7 min per contrast.

#### Map computation and preprocessing

Quantitative MRI maps were computed from the raw MRI data using the hMRI toolbox ^40^ and bespoke analysis scripts written in MATLAB (The MathWorks Inc, Natick, MA, USA). Maps of the MRI parameter R2* were computed separately from the raw echo images with MTw, PDw and T1w contrast, from the regression of the log signal with the corresponding echo times ^48^. The value of the MDI described in Castella et al.^16^ was computed for each R2* map, as provided by the hMRI toolbox^40^. Maps of the MRI parameter MT were computed as described in ^49–53^, following averaging of the echo images for each contrast to increase the signal-to-noise ratio ^54^. The MT maps were only used for spatial normalization of the data into a common group space (see below) and were not used in subsequent analyses.

Data was analysed using Statistical Parametric Mapping (SPM12, Wellcome Centre for Human Neuroimaging, London). The MT maps were segmented into maps of gray and white matter probabilities using Unified Segmentation ^55^. The nonlinear diffeomorphic algorithm Dartel ^56^ was used for inter-subject registration of the tissue classes. The tissue probability maps were normalized to the stereotactic space of the Montreal Neurological Institute (MNI) template using the resulting Dartel template and the deformation fields. As described in Draganski et al.^34^, the quantitative maps were normalized using the same deformation fields but without modulation by the Jacobian determinants. Instead, a combined probability weighting and Gaussian smoothing procedure was used with a 6 mm FWHM isotropic smoothing kernel to produce tissue-specific parameter maps while preserving the quantitative estimates.

#### Analysis

Because the focus of this study was on the methodology involved in incorporating an MDI index into the analysis of MRI data, we restricted our analysis to R2* maps - and MDI values - computed from a single type of raw echo images. We primarily focused on the analysis of the changes in R2* associated with healthy ageing, driven by changes in iron and myelin concentration in grey and white matter respectively ^33,34,57^. Statistical analyses were carried out after estimating the parameters of a general linear model with SPM12. We included four regressors in the model, including age and age^2^, as well as gender and brain volume as variables of no interest. Analyses were conducted using the common approach of assuming identical noise levels in all quantitative maps (*Ordinary Least Squares, OLS*) as well as assuming different noise levels for each map, computed from the MDI values (*Weighted Least Squares, WLS*). The REML algorithm, as implemented in SPM12, was used to estimate a global (spatially invariant) covariance matrix of the data as a linear combination of diagonal matrices built from the MDI values of all participants.

#### Age-associated differences in R2* (age sensitivity)

Statistical F-tests were conducted to assess the significance of age-related differences in R2*. We conducted these analyses, both at the global level of a whole tissue type (grey and white matter) and at the local level of a grey matter region, to assess the performance of WLS analyses in correcting local effects of head motion. The regional analyses were conducted using explicit masks defined from the grey matter maximum probability tissue labels derived from the “MICCAI 2012 Grand Challenge and Workshop on Multi-Atlas Labeling” (https://masi.vuse.vanderbilt.edu/workshop2012/index.php/Challenge_Details), computed from MRI scans originating from the OASIS project (http://www.oasis-brains.org/) and labelled data provided by Neuromorphometrics, Inc. (http://neuromorphometrics.com/) under academic subscription. The regional masks included voxels from both hemispheres.

#### Specificity

We assessed the specificity of the OLS and WLS methods by monitoring the rate of false positives in two types of image analysis frequently conducted in neuroscience studies. I. In a subset of data (N=123; up to 10 images per age bin of 5 years when available), the participants’ age was randomly scrambled between the images before conducting the analysis of age-associated differences in R2* described above. Any positive result would therefore be a false positive. II. In the subset of data with a narrow age range (56-58 years.; N=129), we conducted two-sample t-tests for the comparison of two subgroups. Similarly, any positive results would therefore be a false positive. Analyses were repeated with N_1_ = 2, 5, 10, 20 30 and 60 images in the first group. We repeated both types of specificity analyses 1,000 times, monitoring the rate of false positives across repetitions at the voxel and cluster levels (p<0.05, FWE-corrected). For cluster-level inference, the cluster forming threshold was p<0.001 uncorrected.

#### Accuracy

We investigated the systematic effect of head motion on the R2* estimates (bias) by conducting an F-test analysis of the significance of the dependence of the R2* estimates on the MDI. This was achieved by including regressors in the analysis model, which contained the values of the MDI associated with each image to the power 1 to 4.

## Code availability

Analyses were conducted using custom-made code written in MATLAB R2019b. This code is available here: https://zenodo.org/record/4890217#.YLpB-iVxeUk

The QUIQI method is also available as a plug-in module of the hMRI-toolbox, allowing use of QUIQI from the user interface: https://zenodo.org/record/4737490#.YJJGhB0zaUk. We anticipate merging of this capability with the main version of the toolbox in the near future (https://hmri-group.github.io/hMRI-toolbox/).

## Data availability

Material related to this paper will be made publicly available ^58^. This material consists of data from 123 participants, which include R2* maps computed from raw MTw, T1w and PDw MR images, and grey and white matter tissue probability maps. This data is in the normalized space of the full cohort analysed in this manuscript. This data was selected by binning the participants’ age in bins of 5 years and randomly selecting 10 subjects in each age bin. All subjects were selected in age bins with fewer than 10 subjects.

This material also includes the results of the analyses presented in this manuscript, conducted on the available data. These results can be computed by running the code described in the ‘Code availability’ section on the provided datasets.

This material will be made available to the general public upon publication of this manuscript.

## Acknowledgements

This work was supported by the Swiss National Science Foundation (grant no 320030_184784 (AL)) and the ROGER DE SPOELBERCH Foundation. The Wellcome Centre for Human Neuroimaging is supported by core funding from the Wellcome [203147/Z/16/Z]. CP is Senior Research Associate at the FRS-F.N.R.S., Belgium. The MRI data was acquired on the MRI platform of the Clinical Neuroscience Department, Lausanne University Hospital.

## Author contributions

A.L. designed the experiment. A.L., G.D.D, F.K. and B.D. collected the data. A.L., N.C. and G.D.D. developed techniques and analysed the data. A.L. wrote the paper. A.L., N.C., J.A., G.Z., B.D., C.P., F.K. and M.F.C. designed the analysis and interpreted the results. A.L., N.C., J.A., G.Z., B.D., C.P., F.K., M.F.C. and G.D.D discussed and edited the manuscript.

---

**Extended Data Fig. 1.**
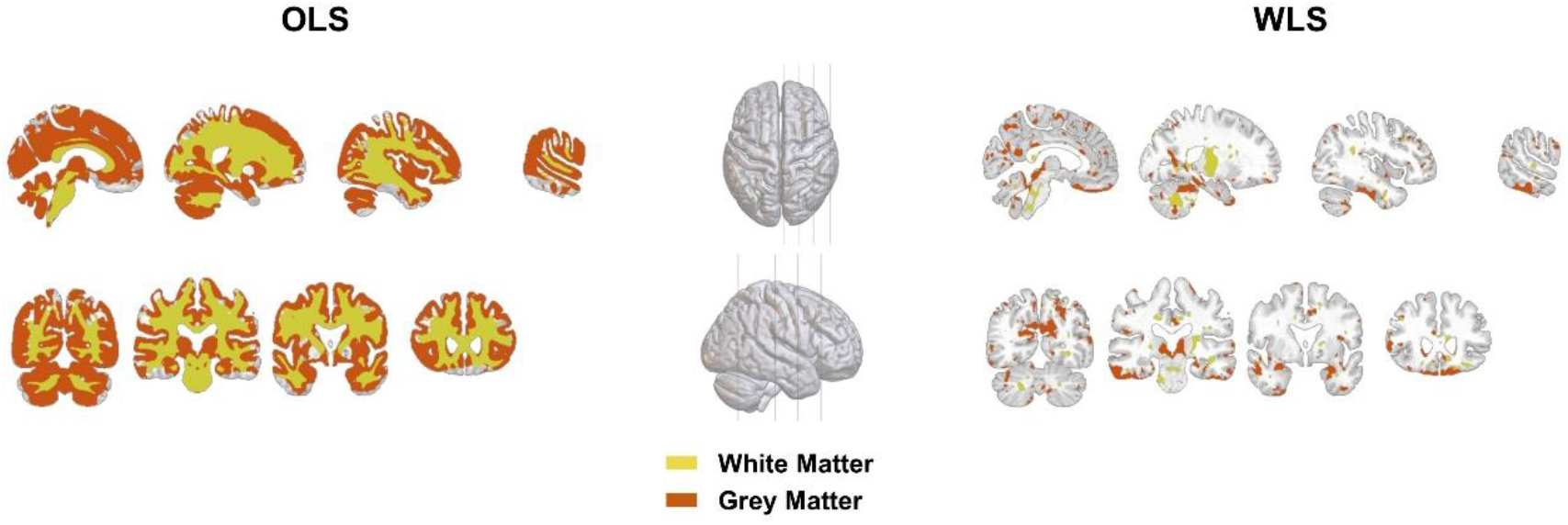
QUIQI reduces residual heteroscedasticity in each voxel of the MRI data. In grey matter, significant heteroscedasticity was found in 82% and 12% of voxels for OLS and WLS respectively (p<0.05). In white matter, significant heteroscedasticity was found in 92% and 9% of voxels for OLS and WLS respectively (p<0.05).

**Extended Data Fig. 2.**
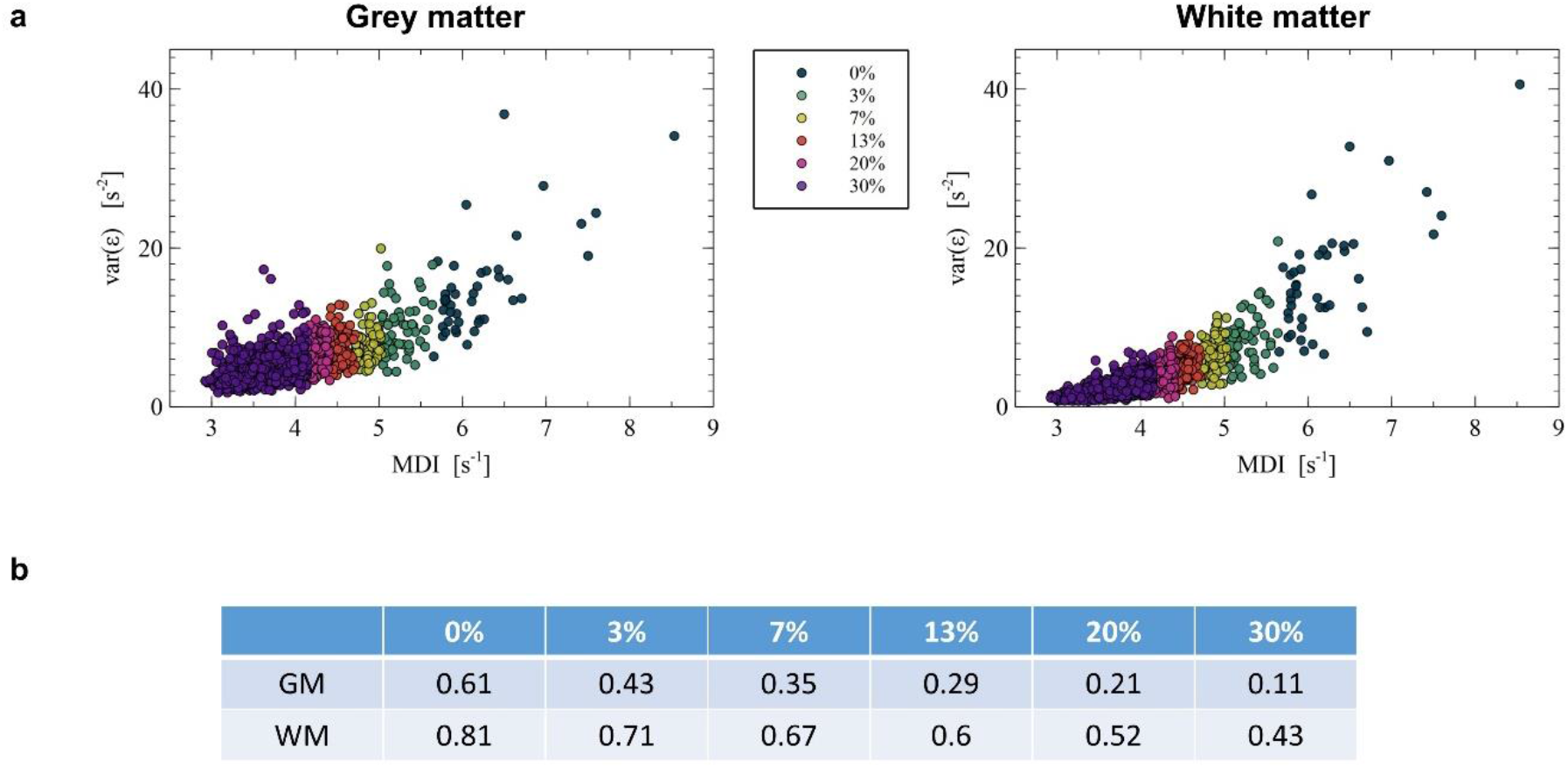
With OLS analyses, restoring homoscedasticity requires removal of 30% of the subjects or more. We assessed noise homoscedasticity after exclusion of up to 30% of the most degraded images from OLS analyses (**a**). We estimated noise homoscedasticity by fitting the dependence of residual noise on the MDI with a polynomial function of order 3. In grey matter, restoring noise homoscedasticity requires the removal of 30% of the images (goodness of fit: R^2^=0.11; (**b**)). In white matter, noise homoscedasticity comparable to that of WLS analyses was not achieved.

**Extended Data Fig. 3.**
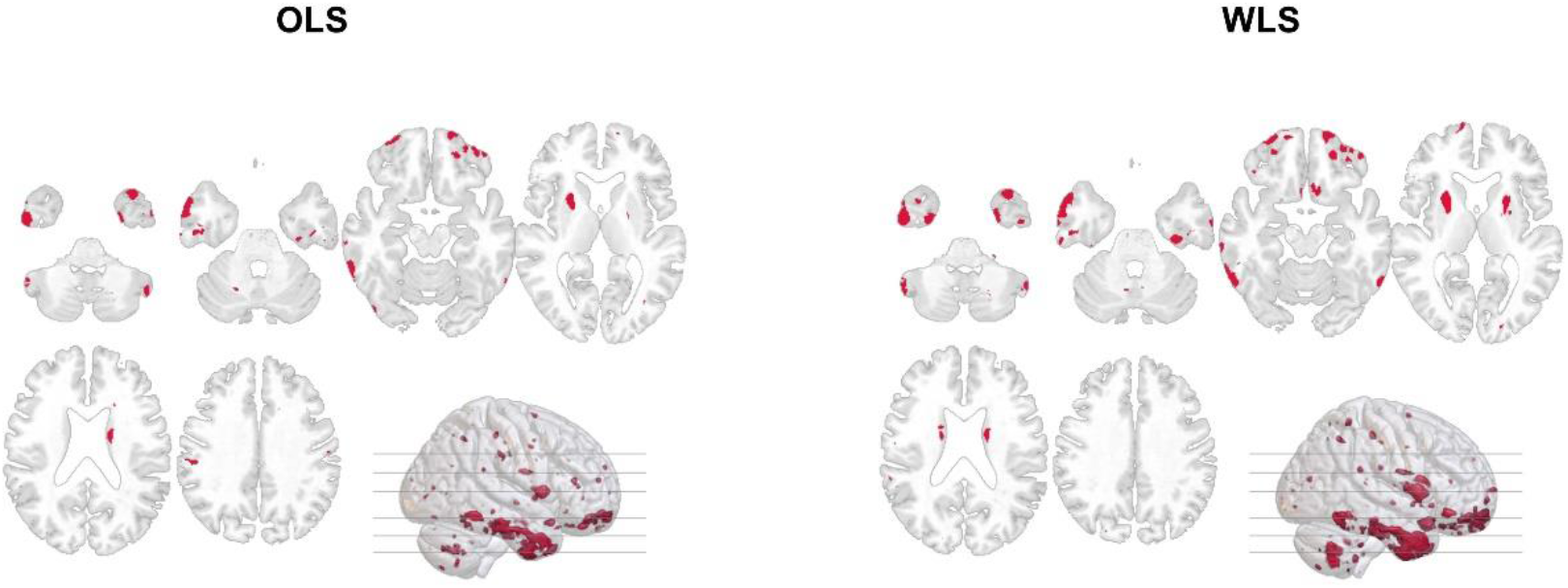
Regions affected by magnetic field inhomogeneities drive the occurrence of false positives in unbalanced group comparisons. Spatial distribution of the voxels showing significant differences at least twice out of 1000 repetitions in the analysis of specificity in group comparisons. The number of subjects in the first group was N1=5. Similarly for OLS (**a**) and WLS (**b**) analyses, these voxels are primarily located in regions affected by magnetic field inhomogeneities (e.g. temporal lobes, orbitofrontal cortex).

**Extended Data Fig. 4.**
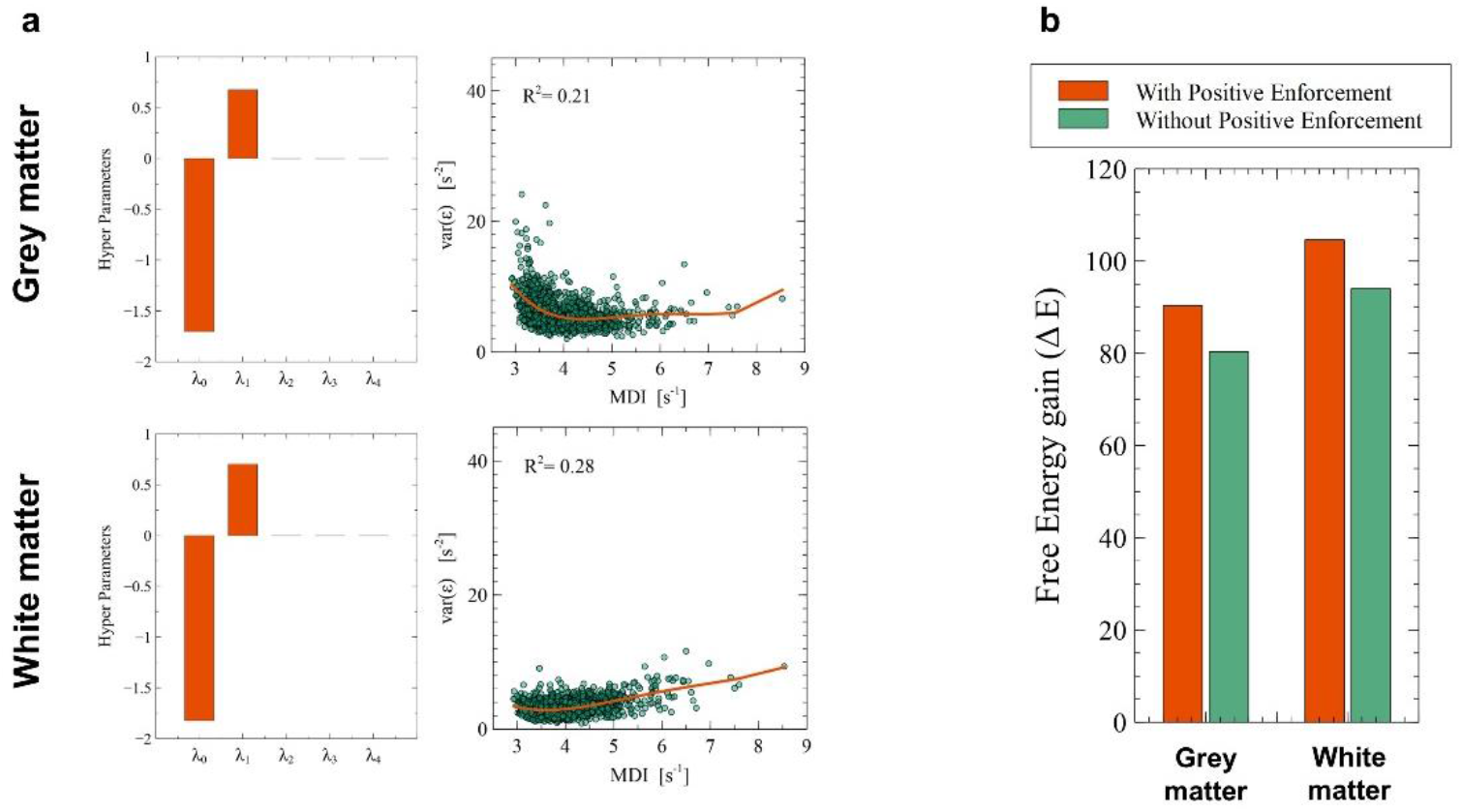
Allowing for negative REML hyper-parameter values leads to a similar performance of QUIQI. The heteroscedasticity of the noise distribution is restored as when positive hyper-parameters is enforced (**a**). Note that consistently with the REML estimation, the polynomial fitting of the dependence of image noise on the MDI also allowed for negative coefficients. The gain in ELBO compared to OLS analyses is also in a similar range (**b**).

**Extended Data Fig. 5.**
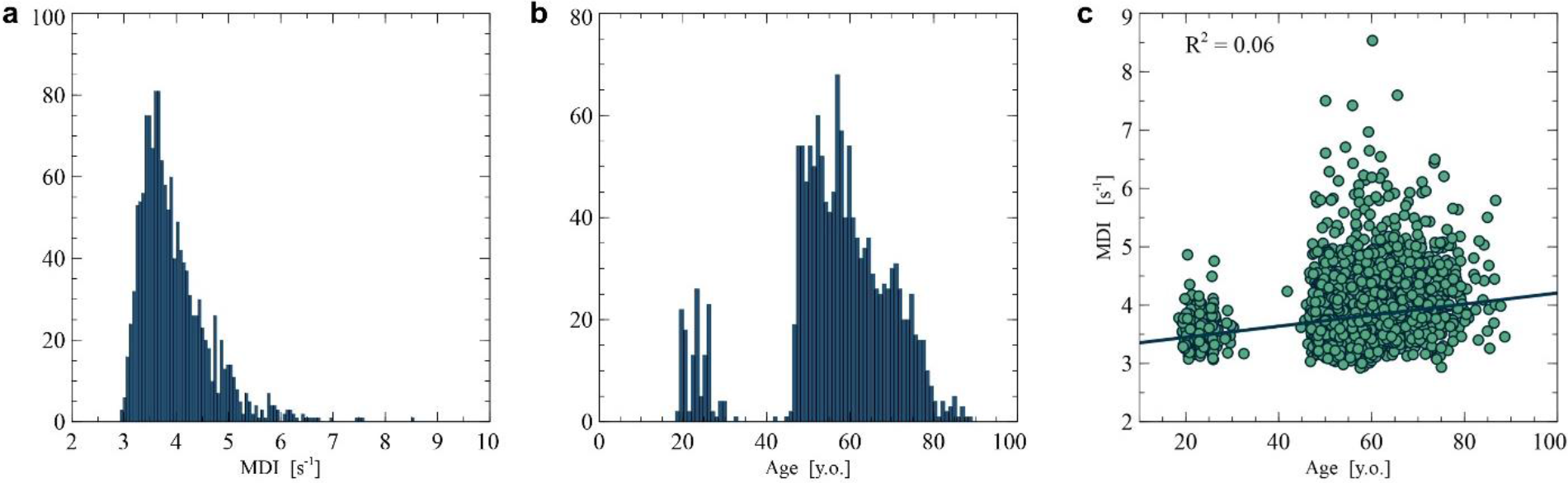
Distribution of the MDI values (a) and participants’ age (b) across the datasets used for analysis (N=1432), and dependence of the MDI on participant’s age (c).

