## Supplementary material for "Restoring statistical validity in group analyses of motion-corrupted MRI data": SIGuide.docx

File title: Supplementary Discussion

This file shows the distribution of MDI values computed from raw MRI data with different contrasts. It also specifies the changes in MDI values after correction for the effects of age and after data normalization. Replicates the main findings from R2* maps computed from T1- and MT-weighted images.
