## Supplementary material for "Restoring statistical validity in group analyses of motion-corrupted MRI data": Supplementaries.pptx

### Slide 1
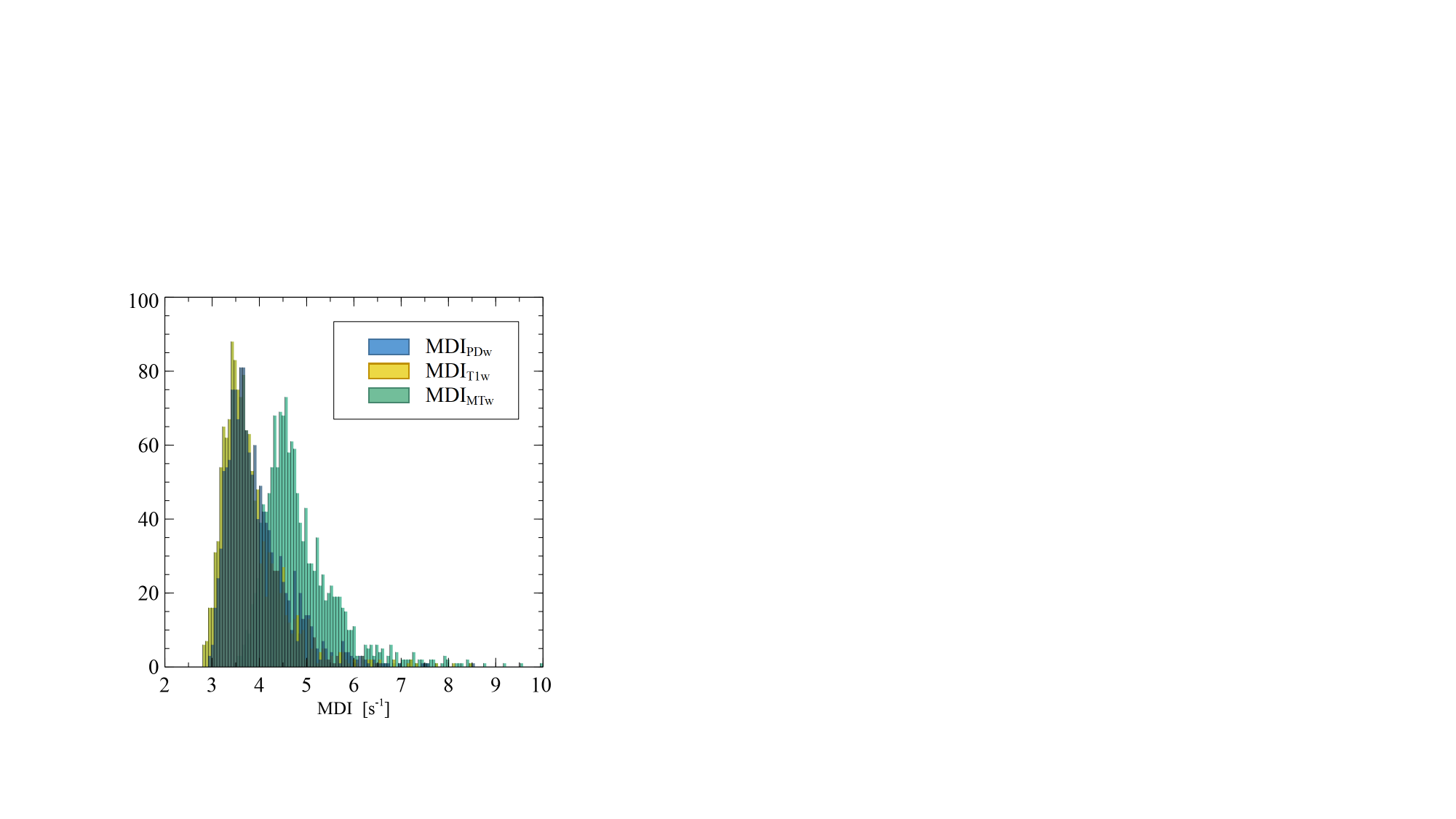

### Slide 2
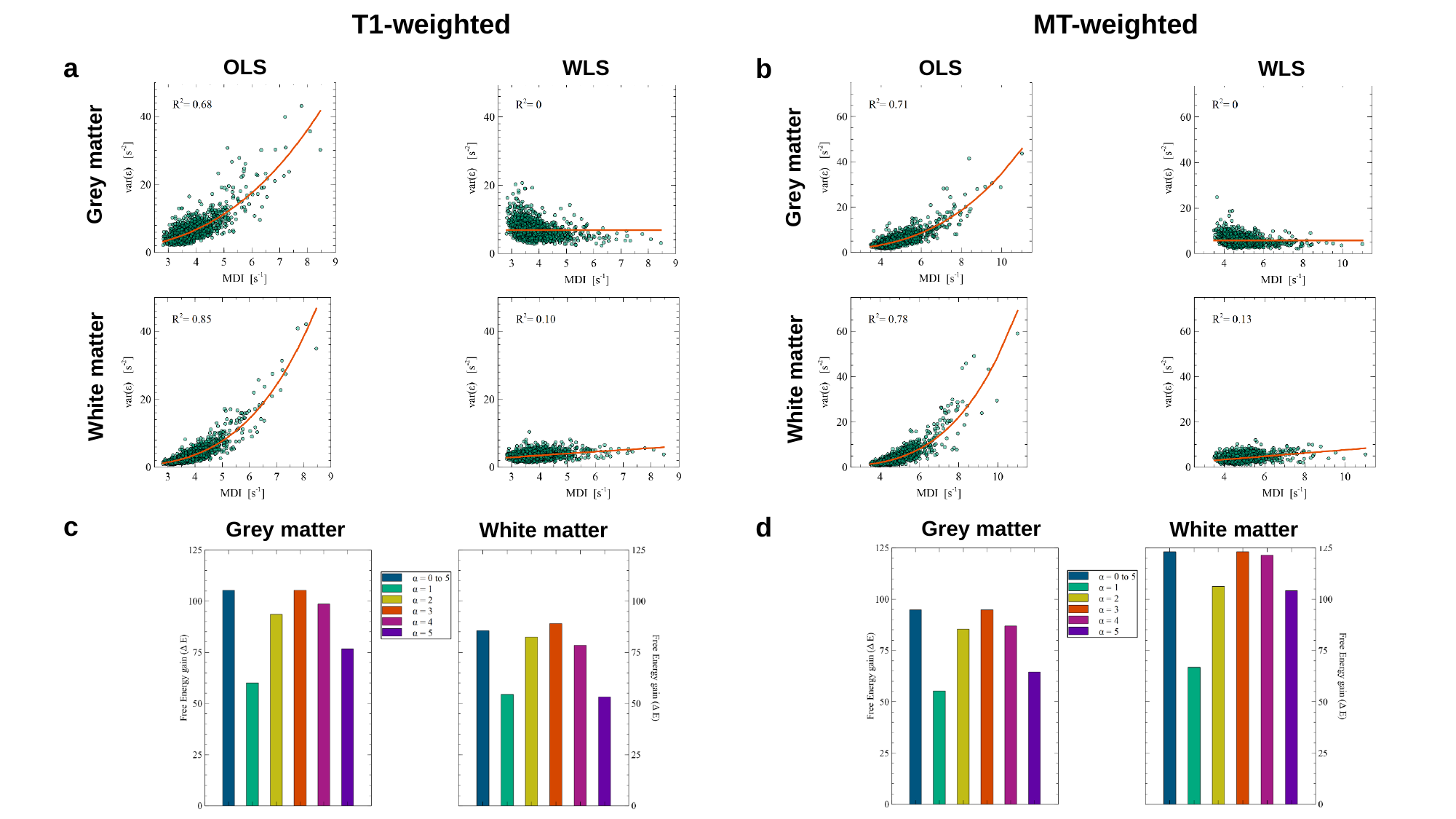

T1-weighted
MT-weighted
a
b
OLS
WLS
OLS
WLS
Grey matter
Grey matter
White matter
White matter
c
d
Grey matter
White matter
Grey matter
White matter
