## Supplementary material for "Restoring statistical validity in group analyses of motion-corrupted MRI data": Supplementary.docx

**Supplementary discussion : confounding factors of the Motion Degradation Index**

Age is the primary source of differences in the BrainLaus MRI data^1^ used here, and affects differentially individual brain regions. In order to estimate the confounding effect of aging on the MDI values, we assessed the changes in MDI values after detrending for age effects. For R2* maps computed from MT-, PD- and T1-weighted raw images, age detrending increased the MDI values by 0.04-0.1s^-1^, small compared to the original MDI values (3-8s^-1^). Also, smoothing after normalization of the data in group space led to a systematic reduction in MDI by 0.2-0,6s^-1^. We also observed a systematic offset in MDI values for R2* maps computed from the MT-weighted images, due to the lower signal-to-noise ratio of the raw data and to the lower number of echo images available for computing the maps (Fig. S1). However, while we illustrated QUIQI using R2* maps computed from PD-weighted images, it performed equally well with R2* maps computed from all image contrasts (Fig. S2). Similarly, we expect QUIQI to show the same level of performance for different image resolutions or number of echo images used for computing the maps.

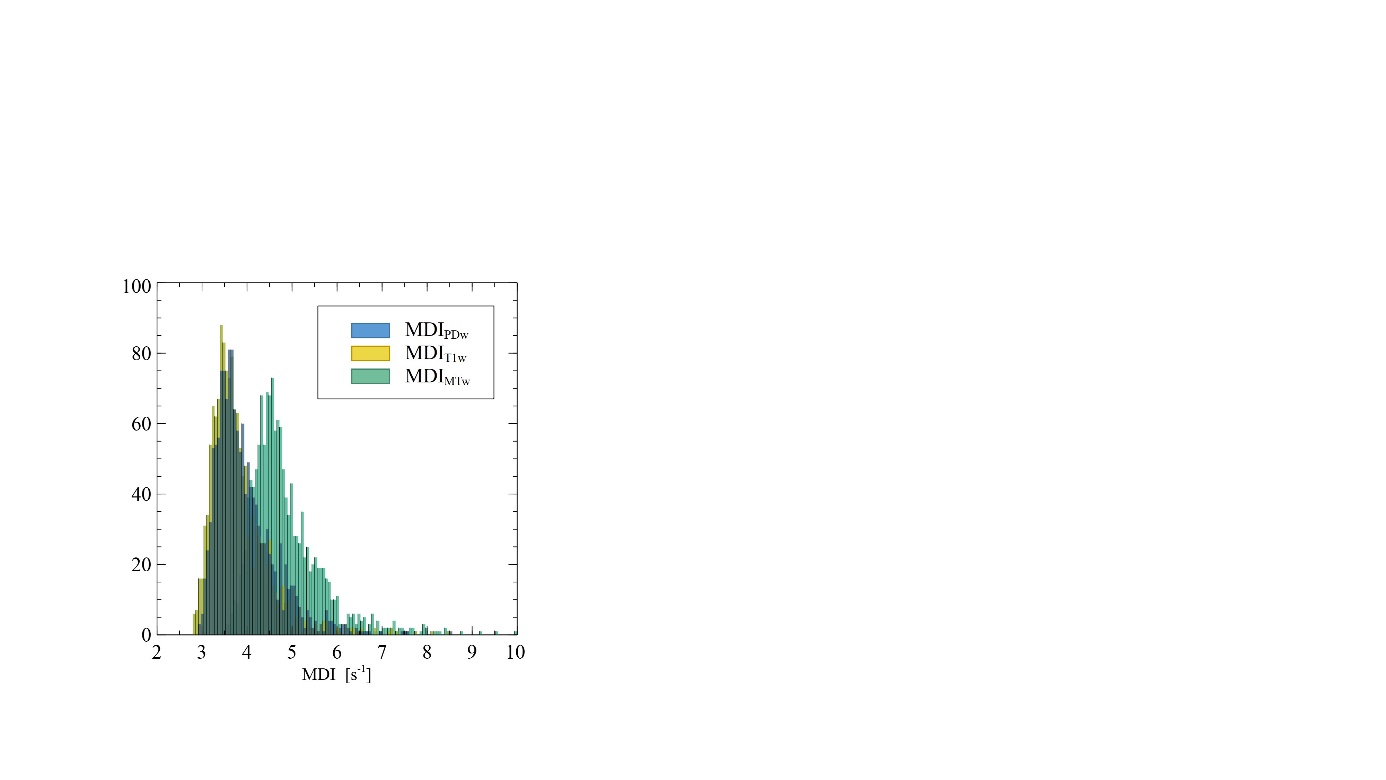

**Supplementary Figure 1. Distribution of the MDI values and of participants’ age across the datasets used for analysis (N=1432).** MRI data with MT-weighted contrast leads to systematically higher MDI values, due to the lower signal-to-noise ratio of the data used to compute this index.

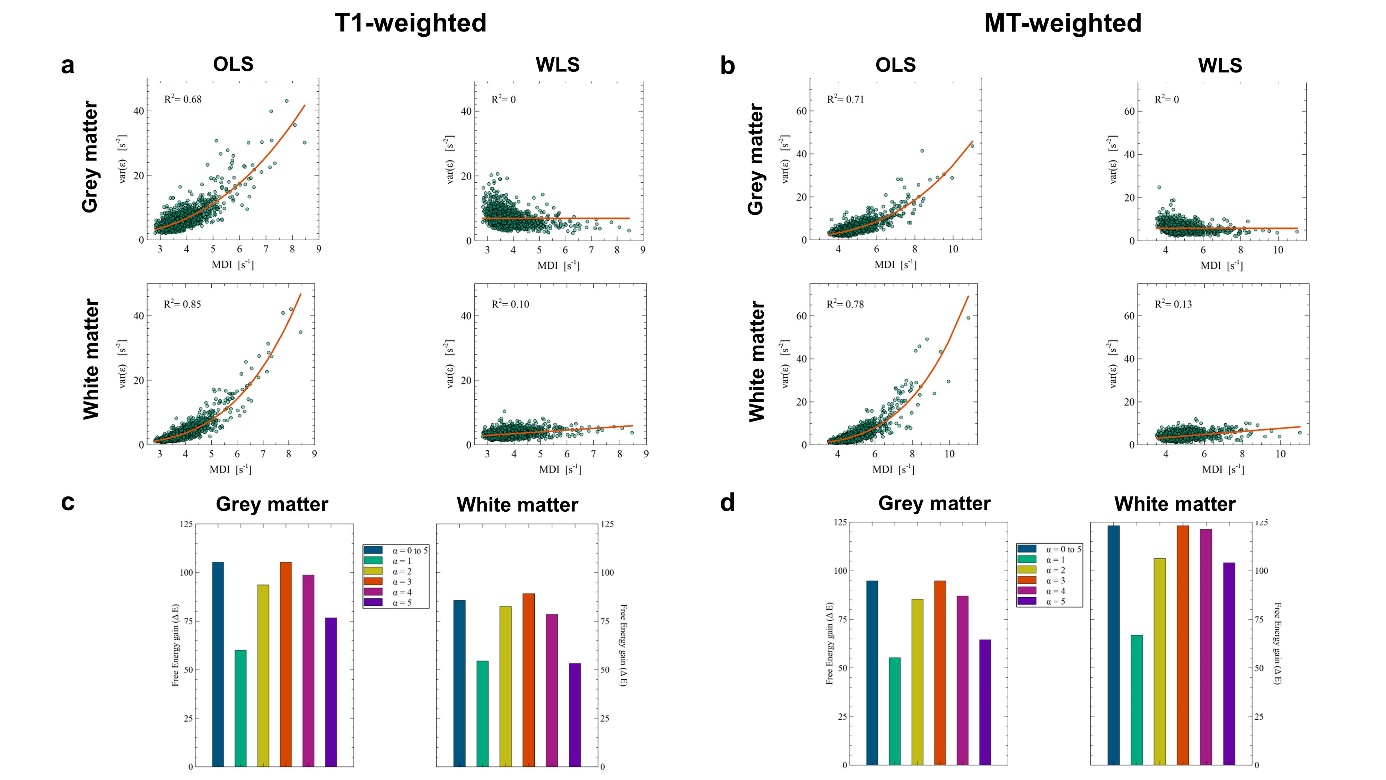

**Supplementary Figure 2. The performance of QUIQI is consistent on R2* maps computed from data with different contrast and signal-to-noise ratio.** QUIQI restores the heteroscedasticity of the noise distribution equally well for T1- (**a**) and MT-weighted data (**b**). The corresponding gain in free energy compared to OLS analyses are consistent across contrasts, and the global optimal noise model also follows a cubic dependence on the motion degradation index (**c**) and (**d**).

1.	Trofimova, O. *et al.* Brain tissue properties link cardio-vascular risk factors, mood and cognitive performance in the CoLaus|PsyCoLaus epidemiological cohort. *Neurobiol. Aging* **102**, 50–63 (2021).
