## Supplementary figures and images for "Restoring statistical validity in group analyses of motion-corrupted MRI data"

### S1.jpg

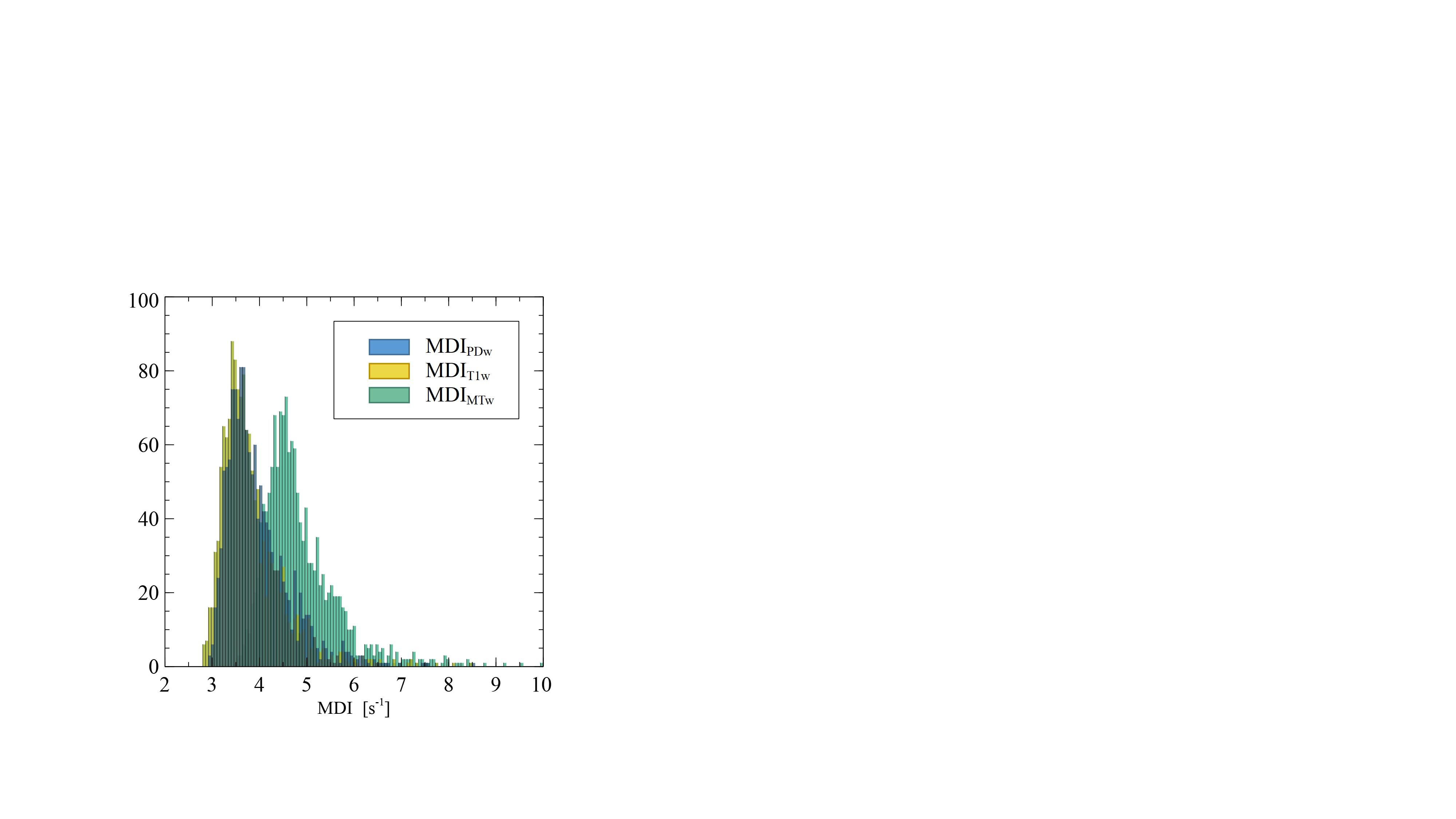

### S2.jpg

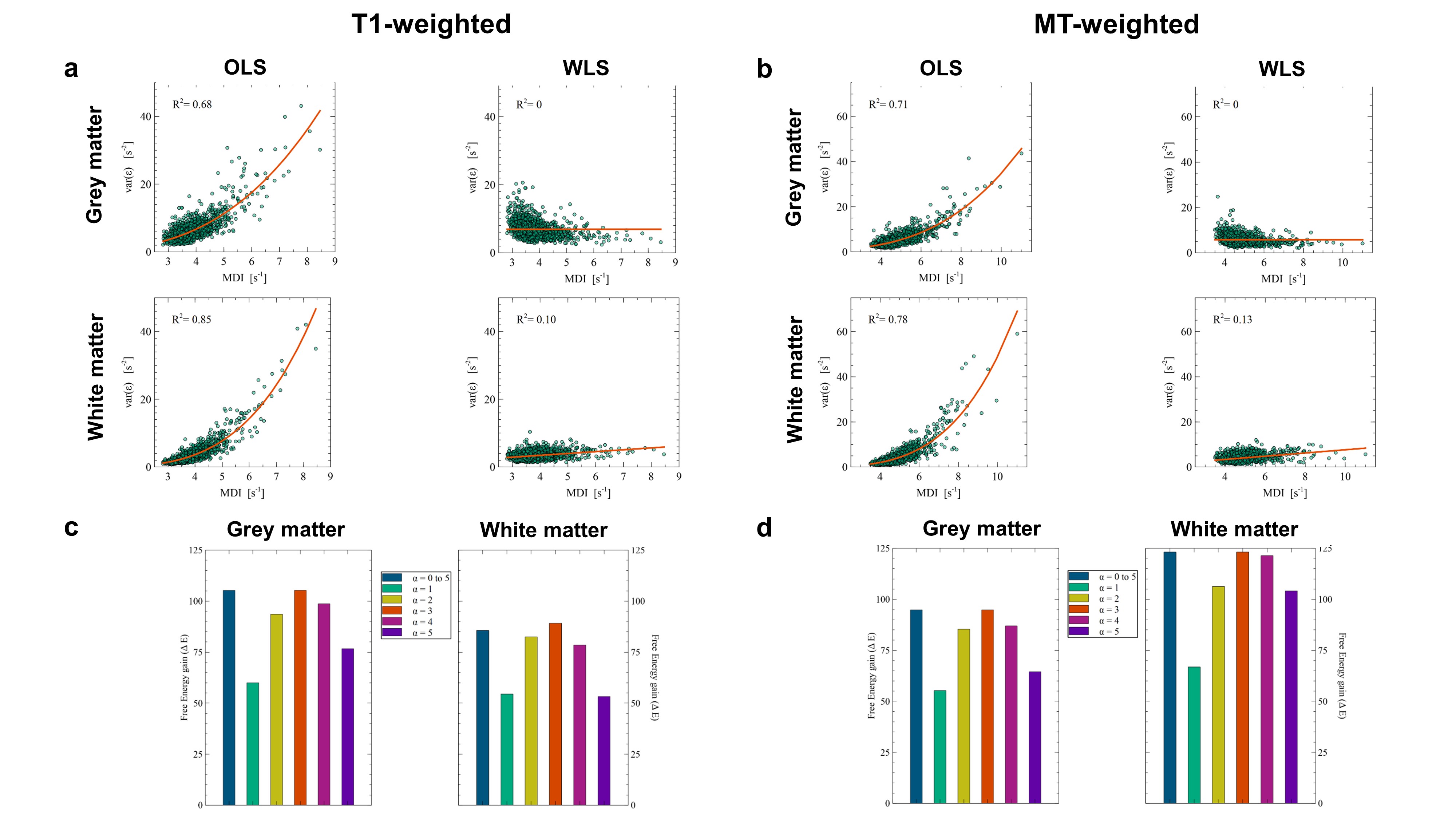
